# Protective effect of agmatine against cisplatin-induced apoptosis in an auditory cell line

**DOI:** 10.1101/2020.03.24.005314

**Authors:** Euyhyun Park, Se Hee Lee, Hoyoung Lee, Young-Chan Kim, Hak Hyun Jung, Gi Jung Im

## Abstract

**Objectives:** Agmatine, an endogenous metabolite of arginine, is known to have antioxidant activity, protect mitochondrial function, and confer resistance to cellular apoptosis. The aim of this study was to evaluate the protective effects of agmatine against cisplatin-induced cellular apoptosis in an auditory cell line.

**Methods:** HEI-OC1 cells were co-treated with agmatine at different concentrations and 15 µM cisplatin for 48 h. Cell viability was measured and annexin V-FITC/propidium iodide (PI) staining was performed to analyze apoptosis. The levels of intracellular reactive oxygen species (ROS) were measured using flow cytometry. The expression of BAX (Bcl2-associated X protein) and the enzymatic activity of caspase-3 were measured to examine the pathway of apoptosis induction.

**Results:** Co-treatment with 8 mM agmatine protected HEI-OC1 cells against cisplatin-induced cell apoptosis. Agmatine exerted a significant protective effect against 15 µM cisplatin when applied for 48 h and reduced the proportion of necrotic and late apoptotic cells. Agmatine did not reduce the cisplatin-induced increase in ROS but decreased the expression of BAX and the activity of caspase-3.

**Conclusions:** Agmatine protected against cisplatin-induced cellular apoptosis in an auditory cell line. These effects were mediated by the protection of mitochondrial function and inhibition of apoptosis.

## Introduction

Cisplatin is one of the most common chemotherapeutic agents used for the treatment of malignant solid tumors [1] as it induces tumor cell death; however, depending on cell type and dose, cisplatin also induces cytotoxicity. Cisplatin-induced cytotoxicity affects the tumor cells as well as other organs; thus, diverse adverse effects may occur, such as neurotoxicity, nephrotoxicity, ototoxicity, and bone marrow suppression [2]. Cisplatin-induced ototoxicity is known to cause hearing loss, tinnitus, ear pain, and dizziness; the extent of this condition is determined by the age of the patients and the cumulative drug dose [3]. It is known that cisplatin induces the production of reactive oxygen species (ROS), which may induce cellular apoptosis in the outer hair cells, spiral ganglion cells, and stria vascularis of the cochlea [4].

Agmatine (decarboxylated arginine) is an endogenous polyamine that is synthesized via the decarboxylation of L-arginine by arginine decarboxylase (ADC) [5, 6]. In previous studies, it has been shown that agmatine has an oxygen radical-scavenging effect and a protective effect on mitochondrial function [7]. In addition, numerous studies have been conducted in the field of neuroscience to examine its effect as a neuromodulator with the function of protecting the CNS in several models of cellular damage [5-9]. Clinically, there is a growing interest in the role of agmatine in epilepsy, Parkinson’s disease, Alzheimer’s disease, depression, spinal cord injury, and neuropathic diseases [10, 11]. However, there have been no reports on the investigation of agmatine in the field of otology.

Consequently, the aim of this study was to evaluate the protective effects of agmatine against cisplatin-induced cellular apoptosis in the HEI-OC1 auditory cell line.

## Materials and Methods

### Chemicals

Agmatine (CAS no. 2482-0-0) and cisplatin (CAS no. 15663-27-1) were obtained from Sigma Aldrich Corporation (St Louis, MO, USA). Agmatine and cisplatin were prepared as 5 mM stock solutions in cell culture medium and diluted to the appropriate concentrations.

### Auditory cell culture

The HEI-OC1 cell line was used as an auditory cell line in this study. The HEI-OC1 cell line was a transgenic immortomouse auditory cell line provided by F. Kalinec (House Ear Institute, Los Angeles, CA, USA). The cells were cultured in high-glucose Dulbecco’s Modified Eagle’s Medium (DMEM; Gibco BRL, Grand Island, NY, USA) containing 10% fetal bovine serum (FBS; JRH Bioscience, Lexena, KS, USA) and 50 U/mL interferon-γ without antibiotics at 33.8 °C and 10% CO_2_ in air. The cell culture used the method of previously reported study [12].

### Cell viability assay

To examine the effect of agmatine on HEI-OC1 cells, we used the WST-based cell viability/cytotoxicity assay kit (EZ-CYTOX, DOGEN, Korea) to measure cell viability in the cell proliferation and cytotoxicity assays. HEI-OC1 cells (2 × 10^4^ cells/well in 48-well microplates) were incubated with 2–20 mM agmatine for 48 h. After incubation, the culture medium was removed and replaced with fresh culture medium; 10 μL of assay solution was added into each well of the microplate, and the microplates were incubated for 2 h at 33.8 °C and 10% CO_2_ in air. The optical density of the contents of each well was measured at 450 nm using a microplate reader (Spectra Max, Molecular Devices, Sunnyvale, CA, USA).

To examine the protective effect of agmatine against cisplatin-induced cellular apoptosis, we co-treated the cells with 2–20 mM agmatine and 15 µM cisplatin (a concentration that was shown to result in a 50% decrease in cell viability in our previous studies) for 48 h. At 48 h after incubation, viability was measured using EZ-CYTOX.

### Flow cytometric assay of apoptosis

Annexin V-fluorescein isothiocyanate (FITC) (Ezway Annexin V-FITC Apoptosis Detection Kit, Komabiotech, Seoul, Korea) was used for determining the pattern of cellular apoptosis. The specific binding of annexin V-FITC occurred during incubation of the cells for 15 min at room temperature in binding buffer containing annexin V-FITC and propidium iodide (PI) at saturating concentrations. A minimum of 10,000 cells was analyzed by FACScan flow cytometry (BD Biosciences, Heidelberg, Germany). The flow cytometric assay used the method of previously reported study [13].

### Intracellular ROS measurement

The fluorescent dye, 2’,7’-dichlorofluorescein diacetate (DCFH-DA, Calbiochem, USA), was used to measure the intracellular ROS level. In the presence of oxidizing agents, DCFH is converted to the highly fluorescent 2’,7’-dichlorofluorescein (DCF). For the analysis, HEI-OC1 cells were incubated in the dark with 50 µm DCFH-DA for 30 min at 37 °C. The FACScan flow cytometer (BD Biosciences, Heidelberg, Germany) was used to analyze the fluorescence at an excitation wavelength of 495 nm and an emission wavelength of 530 nm while gating at 10,000 cells/sample. The measurement of intracellular ROS used the method of previously reported study [13].

### Western blotting analysis of BAX (Bcl-2-associated X protein)

The primary antibodies (BAX (cat no. 2772), beta-actin (cat no. 4967)) were obtained from Cell Signaling Technology (Danvers, MA, USA). Cell lysates were resuspended in radioimmune precipitation assay (RIPA) buffer, and equal amounts of protein (20 µg/sample) were immediately heated at 100°C for 5 min and resolved by sodium dodecyl sulfate polyacrylamide gel electrophoresis (SDS-PAGE). The isolated proteins were transferred to nitrocellulose membranes, and western blotting was performed with gel-loading and protein transfer system kits (Bio-Rad, Hercules, CA). Non-specific binding to the membrane was subsequently blocked by incubation with 5% skim milk in Tris buffer solution containing 0.1% Tween (TBST). The membrane was incubated in solutions of primary polyclonal antibodies at a final dilution of 1:1000. After three washes in TBST, the membranes were incubated with peroxidase-conjugated secondary antibodies in blocking buffer (final dilution, 1:2000) for 1 h and then washed again. A chemiluminescent solution was applied to the membranes (ECL solution; Gendepot, Barker TX, USA), and bound antibodies were detected using a ChemiDOC touch imaging system (Bio-Rad, Hercules, CA). The relative band densities were computed using ImageJ software (Imagej.nih.gov/ij/index.html).

### Measurement of caspase-3 activity

The enzymatic activity of caspase-3 was analyzed using a caspase3/CPP32 fluorometric assay kit (Biovision, Milpitas, California, USA). HEI-OC1 cell lysates were prepared in lysis buffer on ice for 10 min and centrifuged at 14,000 rpm for 5 min. The protein concentration of each lysate was measured. The catalytic activity of caspase-3 in the cell lysates was determined by measuring the proteolytic cleavage of 50 mM DEVD-AFC and fluorometric substrates at 37°C for 2 h. Cell lysate mixture not incubated with DEVD-AFC as a substrate was used as a negative control. The plates were analyzed using a microplate reader with a 400 nm excitation filter and a 505 nm emission filter (Spectra Max, Molecular Devices, Sunnyvale, CA, USA). The measurement of caspase-3 activity used the method of previously reported study [13].

### Statistical analysis

All values are presented as the mean ± standard deviation (SD) using the SPSS 22.0 statistical program. The paired *t*-test was used to analyze data pairs, and P values < 0.05 were considered statistically significant.

## Results

### Effect of agmatine on HEI-OC1 cells and protective effect of agmatine against cisplatin-induced cytotoxicity

HEI-OC1 cells were co-treated with 2–20 mM agmatine for 48 h and cell viability was measured using the EZ-CYTOX assay. The maximal protective effect was observed for 10 mM agmatine, with significant protective effects compared with the control found with 8 and 10 mM agmatine (***P<0.001, respectively). Agmatine concentrations above 20 mM resulted in lower cell viability than the control. The results were obtained from five separate experiments performed in triplicate (Fig. 1A).

**Fig 1.**
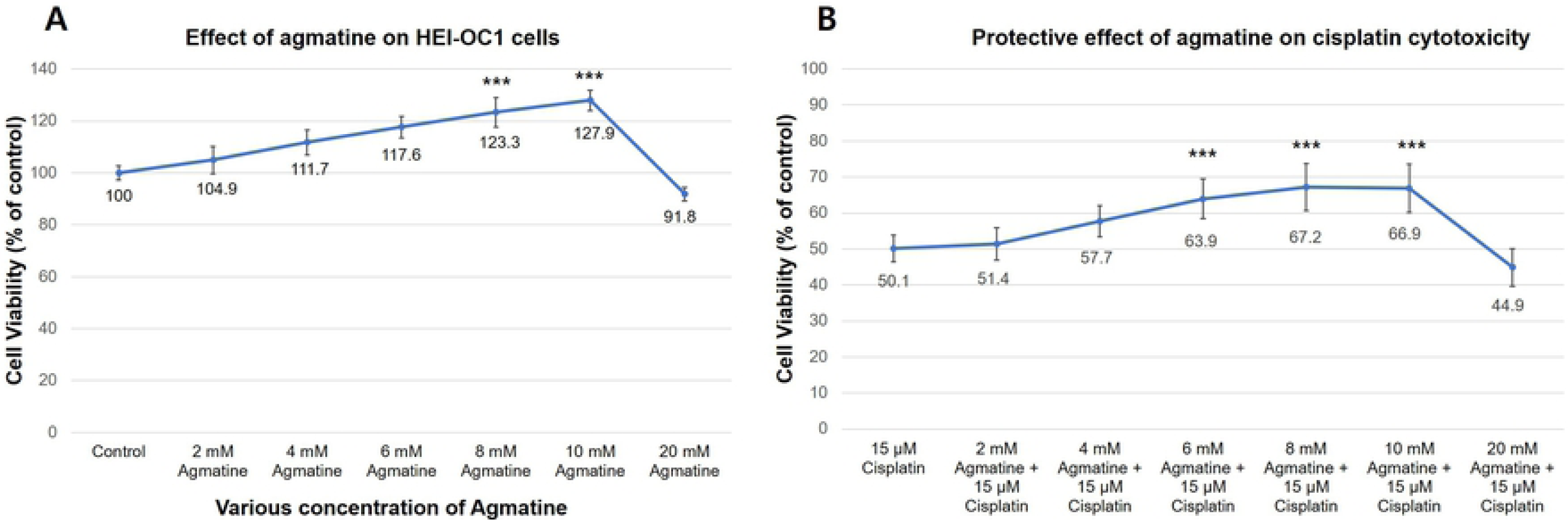
Effect of agmatine on HEI-OC1 cells. (A) HEI-OC1 cells were co-treated with agmatine (AG; 2–20 mM) for 48 h. The maximal protective effect of agmatine was observed at a concentration of 10 mM, and a significant protective effect was observed at 8 and 10 mM of AG compared with the control (***P < 0.001). (B) HEI-OC1 cells were co-treated with 2–20 mM AG concentrations and 15 µM cisplatin for 48 h. AG (6–10 mM) provided significant protection against cytotoxicity induced by 15 µM cisplatin (vs. 50.1% ± 3.7% viability in the cisplatin group). In normal conditions, the maximal protective effect was observed from 10 mM AG. However, in the presence of cisplatin, the maximal protective effect of AG was observed at a concentration of 8 mM. The results were obtained from five separate experiments performed in triplicate (***P < 0.001).

The protective effect of agmatine on cisplatin-induced cytotoxicity in HEI-OC1 cells was measured. HEI-OC1 cells were co-treated with variable concentrations of agmatine (2–10 mM) with 15 µM cisplatin for 48 h. Agmatine (6–10 mM) provided significant protection against the cytotoxic effects of 15 µM cisplatin in the (vs. 50.1% ± 3.7% viability in the cisplatin group, ***P < 0.001, respectively). In normal conditions, the maximal protective effect occurred with 10 mM agmatine. However, in the presence of cisplatin, the maximal protective effect was observed from 8 mM agmatine. Thus, 8 mM was chosen as the ideal agmatine concentration for analysis of the protective effects against cisplatin-induced cytotoxicity. The results were obtained from five separate experiments, each performed in triplicate (Fig. 1B).

Representative microscopy images show the protective effect of agmatine against cisplatin-induced cytotoxicity in HEI-OC1 cells. In the 8 mM agmatine group, slight cellular growth was observed, similar to the control group (Fig. 2A and 2B). In the 15 µM cisplatin group, significant necrotic debris and a decreased cell population was observed on the surface of the cell culture (Fig. 2C). Co-treatment with 8 mM agmatine resulted in cell sizes within the normal range and a decrease in necrotic debris compared with the cisplatin group (Fig. 2D).

**Fig 2.**
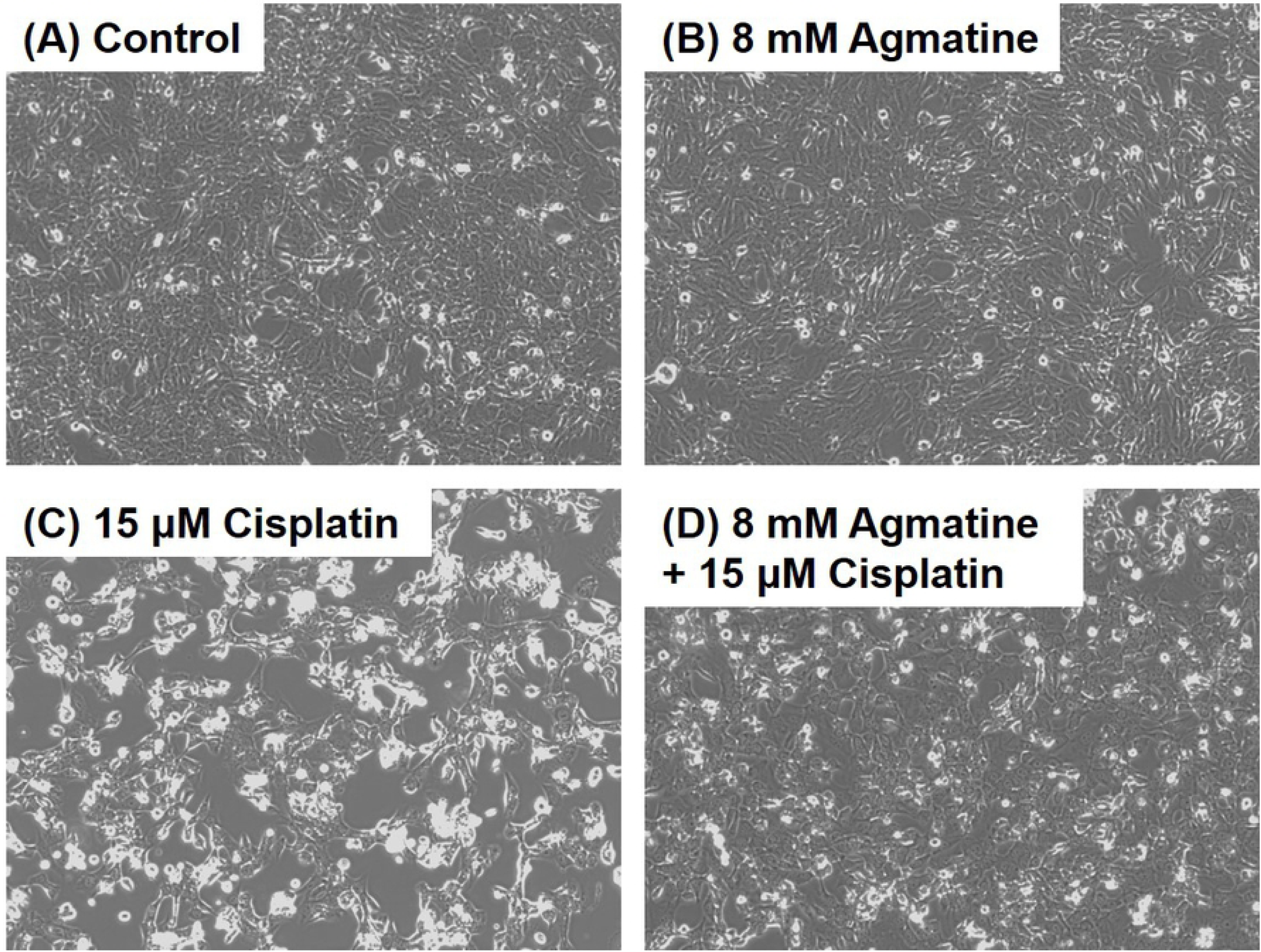
Representative microscopy images showing the protective effects of agmatine against cisplatin-induced cytotoxicity in cultured auditory cells. (A) Control. (B) 8 mM Agmatine. Agmatine promoted slight cellular growth similar to the control group. (C) 15 µM Cisplatin. Significant necrotic debris and a decrease in the cell population were observed on the cell culture surface. (D) Cisplatin + Agmatine group. Agmatine prevented cisplatin-induced cytotoxicity; cell sizes were within the normal range and necrotic debris was lower than that in the cisplatin group.

### Agmatine significantly reduced the proportion of cells in necrosis and late apoptosis induced by cisplatin

Flow cytometric analysis was used to identify the necrotic or late apoptotic cells (Fig. 3A). In both the control and 8 mM agmatine groups, the percentages of necrotic and late apoptotic cells were similar (2.2% ± 1.1% and 5.3% ± 1.3% in the control group; 3.4% ± 2.7% and 8.3% ± 3.9% in the agmatine group). In the 15 µm cisplatin group, significantly higher density was observed for both necrosis and late apoptosis area (9.2% ± 3.0% and 33.5 ± 5.6%, respectively), compared with that in the control and agmatine groups (*P < 0.05). Agmatine treatment significantly reduced both necrosis and late apoptosis (Fig. 3B): necrosis was reduced from 9.2% to 4.8% ± 1.7% (*P < 0.05) and late apoptosis was reduced from 33.5% to 21.3% ± 4.9% (**P < 0.05). The results were obtained from five separate experiments.

**Fig 3.**
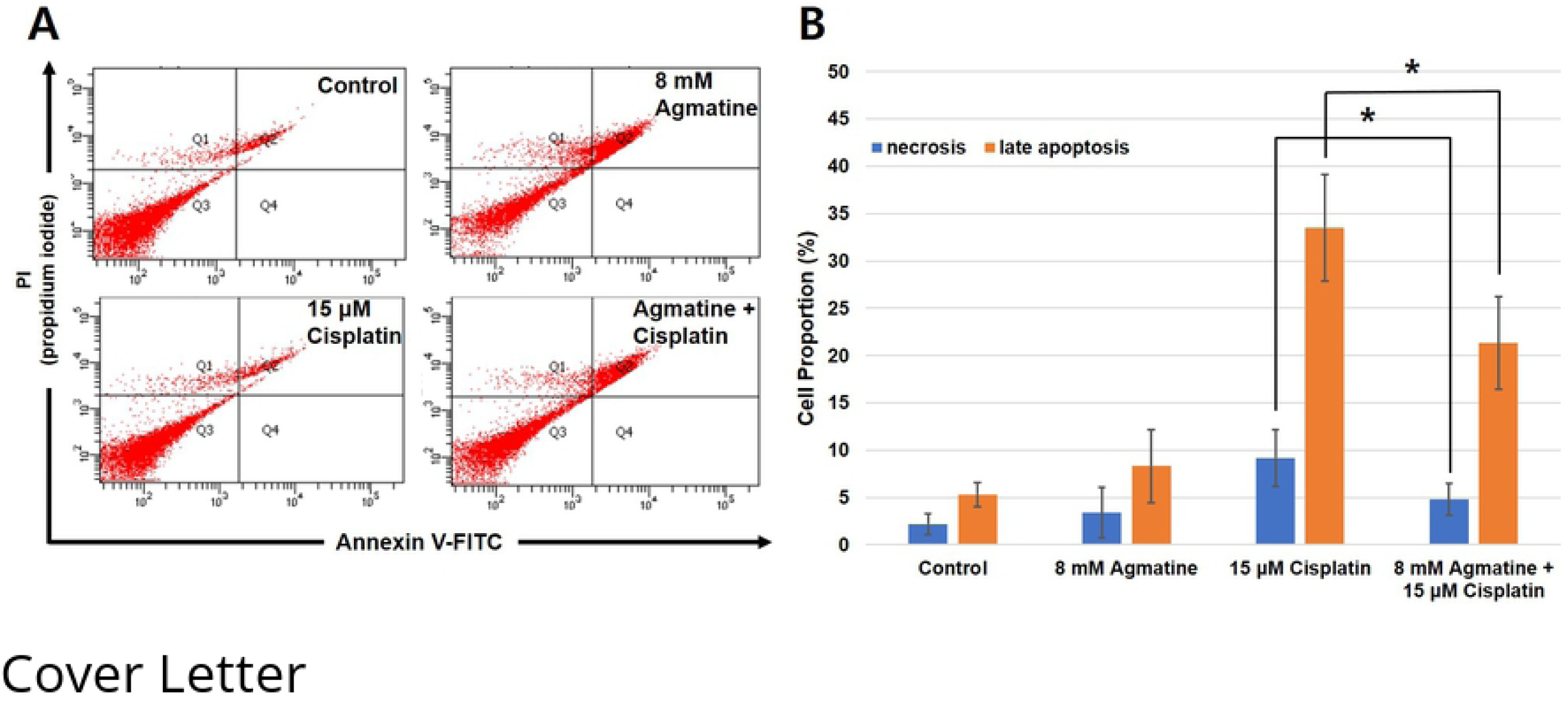
Apoptosis study of agmatine. (A) Representative data for flow cytometric analysis showing cellular apoptosis patterns of necrosis (left-upper panel) or late apoptosis (right-upper panel). In the 15 µM cisplatin group, an increase in the percentage of cells in late apoptosis was observed. However, agmatine treatment reduced the density in both upper panels. (B) The application of 15 µM cisplatin induced a significant increase in the percentages of cells in necrosis and late apoptosis, compared with the control and agmatine groups (**P < 0.05). Agmatine treatment significantly reduced both necrosis and late apoptosis. Necrosis was reduced from 9.2% to 4.8% ± 1.7% (*P < 0.05), and late apoptosis was reduced from 33.5% to 21.3% ± 4.9% (**P < 0.05). The results were from five separate experiments.

### Agmatine did not reduce the cisplatin-induced increase in ROS

To investigate the effects of agmatine on the intracellular ROS generation induced by cisplatin, HEI-OC1 cells were treated with 15 µM cisplatin in the presence or absence of 8 mM agmatine for 48 h. Compared with the control group, the cisplatin group showed a significant increase in ROS generation (2.75 ± 0.27fold, ***P < 0.001). However, 8 mM agmatine treatment did not significantly reduce ROS (2.56 ± 0.28-fold vs 2.75 ± 0.27-fold increases, respectively, compared with the control group; P = 0.3 for the agmatine vs. cisplatin comparison). The results were obtained from five separate experiments.

### Agmatine inhibited the expression of BAX related to cisplatin-induced apoptosis

BAX is a known pro-apoptotic protein in the mitochondrial apoptosis pathway. In the cisplatin group, increased activity of BAX was observed compared with the control group (***P < 0.001). The co-treatment of HEI-OC1 cells with 8 mM agmatine and cisplatin significantly decreased the activity of BAX compared with cisplatin treatment alone (*p < 0.05) (Fig. 4). The results were obtained from five separate experiments.

**Fig 4.**
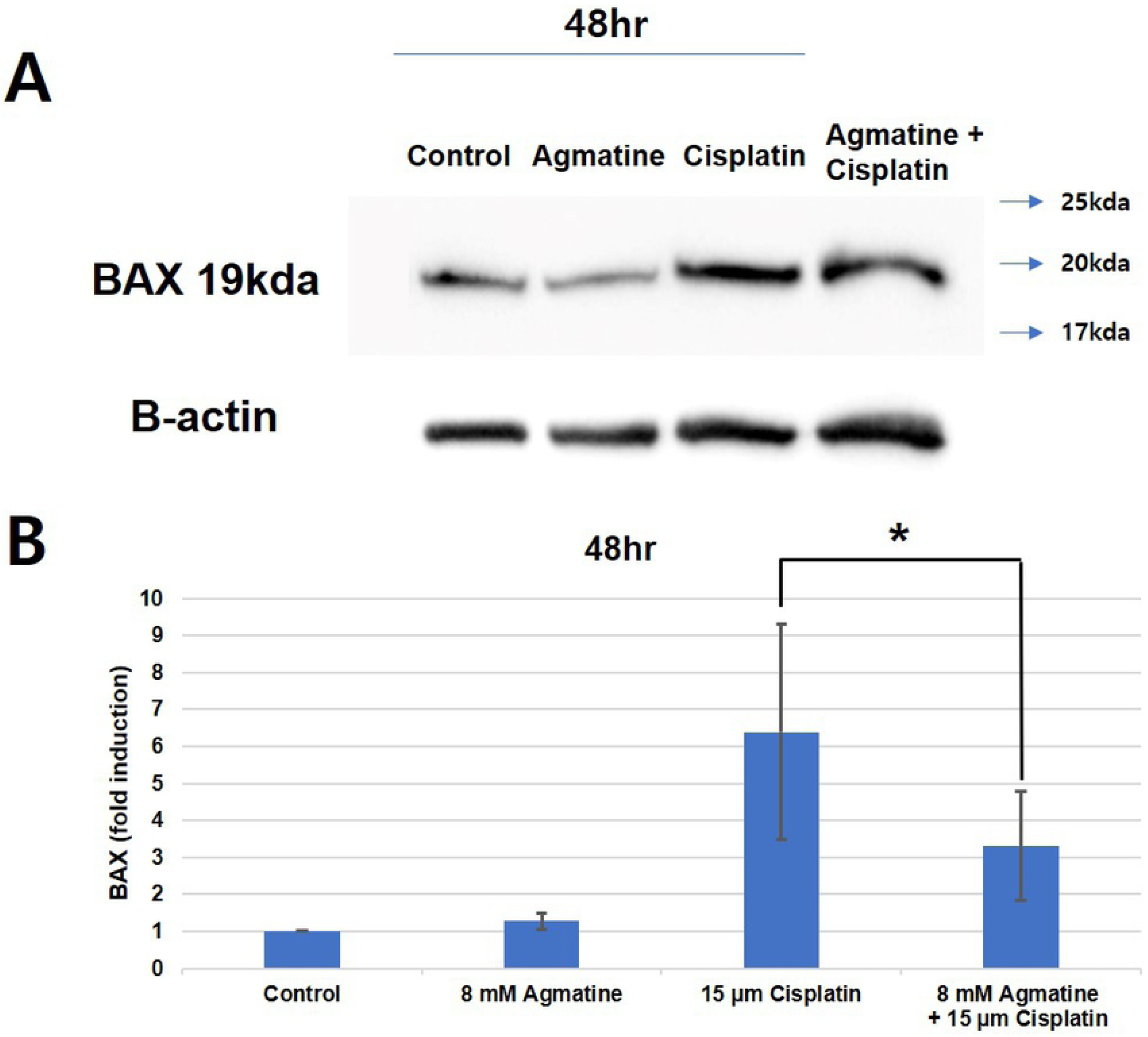
Western blotting of BAX. In the cisplatin group, increased activity of BAX was observed. The activity of BAX in the HEI-OC1 cells co-treated with 8 mM agmatine was significantly lower than that in the cells treated with cisplatin alone (*p < 0.05). The results were obtained from five separate experiments.

### Agmatine inhibited the expression of caspase-3 activity

Caspase-3 activity is involved in cisplatin-induced toxicity and is related to the apoptotic changes that occur in cisplatin ototoxicity. The administration of 15 µ M cisplatin increased the activity of caspase-3 (16.04 ± 1.74-fold compared with control cells). Co-treatment of HEI-OC1 cells with 8 mM agmatine and cisplatin significantly reduced caspase-3 activity (9.68 ± 0.78-fold compared with the normal control) compared with cells treated with cisplatin alone (***P < 0.001) (Fig. 5). The results were obtained from five separate experiments.

**Fig 5.**
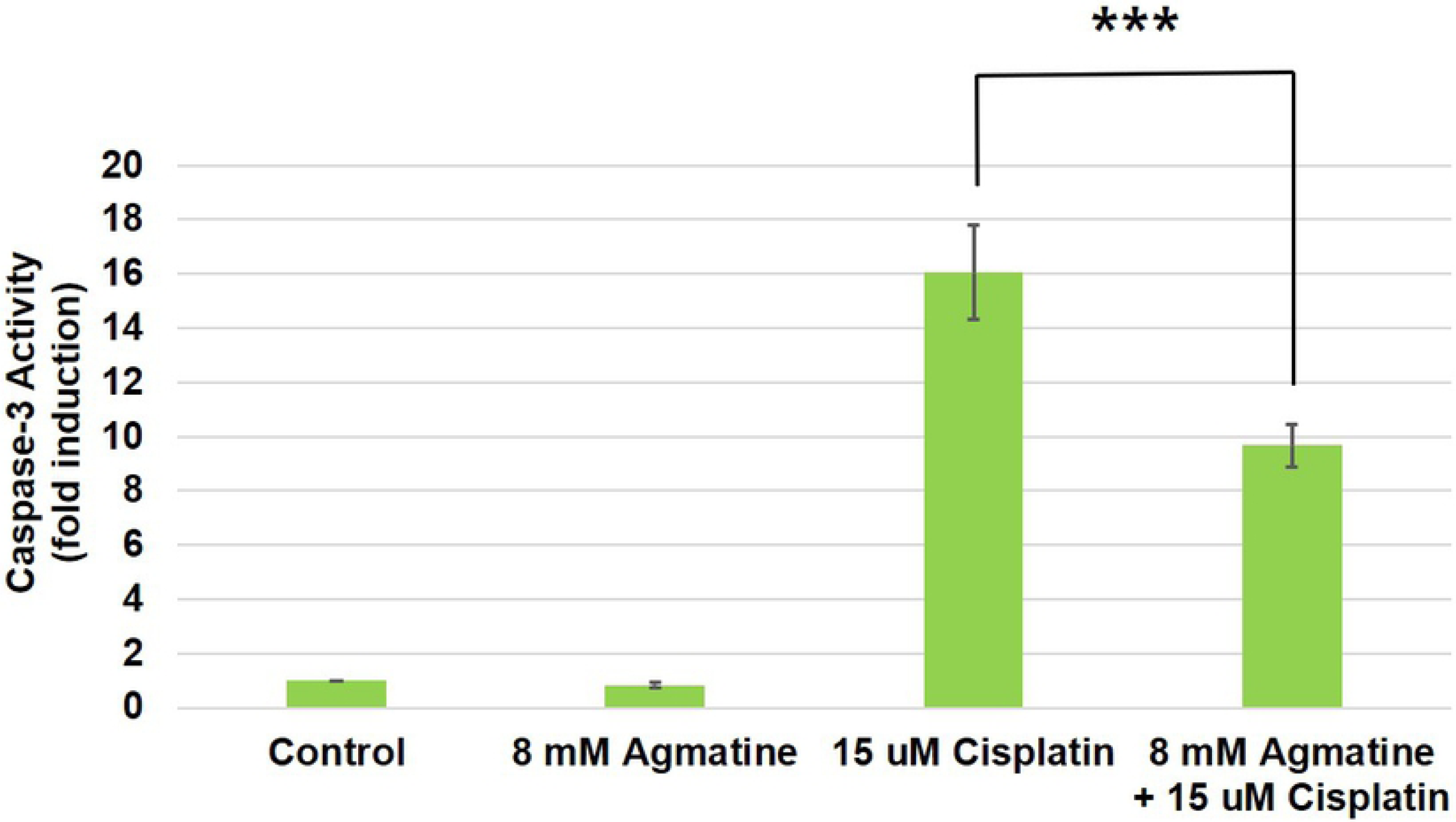
Measurement of caspase-3 activity. The data showed that co-treatment with agmatine, compared with cisplatin alone, inhibited the expression of caspase-3 (16.04 ± 1.74-fold compared with the normal control for the cisplatin group vs. 9.68 ± 0.78-fold compared with the normal control for the cisplatin-plus agmatine group) (***p < 0.001, compared with the cisplatin-treated group). The results were obtained from five separate experiments.

## Discussion

Agmatine, an endogenous metabolite of arginine, is known to have antioxidant activity, protect mitochondrial function, and confer resistance to cellular apoptosis [7]. In this study, agmatine significantly protected against cisplatin-induced apoptosis in HEI-OC1 cells. Agmatine did not significantly reduce ROS but was found to inhibit cellular apoptosis via protective effects on the mitochondrial function of HEI-OC1 cells in this study. To date, this is the first study, to our knowledge, to investigate the protective effects of agmatine on auditory cells.

It is well known that cisplatin causes potentially irreversible damage to auditory hair cells; clinically, this may lead to sensorineural hearing loss and tinnitus [14]. The sensorineural hearing loss appears to be dose related, cumulative, bilateral, and usually permanent and initially occurs in the higher frequencies [15, 16]. A known mechanism of cisplatin ototoxicity is the generation of excessive ROS by cisplatin. ROS can deplete antioxidant enzymes in the cochlear tissue and can increase calcium influx and apoptosis in hair cells of the cochlea [15]. ROS activates BAX in the cytosol, translocating it to the mitochondria and allowing cytochrome c release into the cytosol; it can also activate caspase-3 and -9, which induces cellular apoptosis [17, 18]. As mentioned above, agmatine did not reduce the excessive ROS induced by cisplatin in this study, but it was considered to protect auditory cells from apoptosis through a reduction in the expression of BAX.

Song et al. reported that diphenyleneiodonium reduced cellular apoptosis in kidney proximal tubular epithelial cells without a reduction in ROS owing to the upregulation of Bcl2 [19]. Cellular apoptosis was reduced by maintaining mitochondrial membrane permeabilization through Bcl2 family regulation; the results of our present study are considered to be similar to those of that study.

Agmatine is a natural product that was discovered more than 100 years ago; however, research on this compound is still in progress. The known clinical effects include neuroprotection, neuropathic pain reduction, nerve regeneration, and antidepressant and antioxidant effects [20]. Therefore, although agmatine is currently a compound of interest in the field of neuroscience, its potential activities related to otology have not yet been explored. In previous studies, agmatine was shown exert to protective effects against cell damage and cellular apoptosis in human neuroblastoma cells, hippocampal neuronal cells, and retinal neuronal cells [5-9, 21, 22]. Although this study investigated the protective effects of agmatine on auditory cells, previous studies have mainly shown its protective effects on neuronal cells. Thus, the authors expect that the protective effect of agmatine on spiral ganglion neuronal cell may be determined in future studies.

The authors conducted this study to investigate if the action of agmatine as an N-methyl-D-aspartate (NMDA) receptor antagonist would be effective for the treatment of subjective tinnitus. Currently, numerous studies have examined on the effect of NMDA receptor antagonist in tinnitus treatment [23]. In addition, known effects of agmatine, such as pain reduction, nerve regeneration, and antidepressant effects, are expected to improve tinnitus. In our experiments performed on an auditory cell line, it was confirmed that agmatine protected auditory cells from cisplatin-induced cytotoxicity and did not result in ototoxicity. In future, we intend to investigate if agmatine is an effective tinnitus therapy and if it protects against auditory cells in ex vivo and in vitro studies.

## Conclusions

Agmatine exerted significant protective effects against cisplatin-induced cellular apoptosis in HEI-OC1 auditory cells. These effects were mediated by the protection of mitochondrial function and the inhibition of apoptosis.

